# Disease-associated Kv1.3 variants are energy compromised with impaired nascent chain folding

**DOI:** 10.1101/2025.01.17.631970

**Authors:** Aaron Sykes, Lannawill Caruth, Shefali Setia Verma, Toshinori Hoshi, Carol Deutsch

## Abstract

Human Kv1.3, encoded by *KCNA3*, is expressed in neuronal and immune cells. Its impaired expression or function produces chronic inflammatory disease and autoimmune disorders, the severity of which correlates with Kv1.3 protein expression. The intersubunit recognition domain, T1, at the cytosolic N-terminus of Kv1.3, acquires secondary, tertiary, and quaternary structures during early biogenesis while the nascent protein is attached to the ribosome and/or the ER membrane. In this study, we ask whether native *KCNA3* gene variants in T1 are associated with human disease and whether they manifest early-stage folding defects, energetic instabilities, and conformational distortion of subunits. We use three approaches: first, the unbiased “genome-first” approach to determine phenotype associations of specific *KCNA3* rare variants. Second, we use biochemical assays to assess early-stage tertiary and quaternary folding and membrane association of these variants during early biogenesis. Third, we use all-atom molecular dynamics simulations of the T1 tetramer to assess structural macroscopic and energetic stability differences between wildtype (WT) Kv1.3 and a single-point variant, R114G. Measured folding probabilities and membrane associations are dramatically reduced in several of the native variants compared to WT. Simulations strikingly show that the R114G variant produces more energetically unstable and dynamic T1 domains, concomitant with tertiary unwinding and impaired formation of symmetrical tetramers. Our findings identify molecular mechanisms by which rare variants influence channel assembly, potentially contributing to diverse clinical phenotypes underlying human disease.

## INTRODUCTION

The physicochemical processes by which a mature ion channel forms remain largely unknown. During translation of channel mRNA, a stepwise progression of protein folding events occurs, including the coupled acquisition of secondary, tertiary, and quaternary conformations and induced physicochemical interactions. These successive folding steps early in the process profoundly affect the final protein structure^1,2^. Even a single mutation at a critical site can impair proper folding. Such folding disruptions often lead to pathological or lethal outcomes^3–5^, e.g., cardiovascular disease, cancer, immunological and neurological disorders^6^. This deleterious scenario is particularly striking for voltage-gated K^+^ (Kv) channels (Fig. 1A), which are present across all biological kingdoms and absolutely essential for the normal physiological function of both excitable and nonexcitable cells^7,8^. Misfolding of Kv channels is implicated in a diverse array of disorders, however, an extensive gap exists in our mechanistic understanding of this process, severely limiting development of effective strategies for therapeutic intervention. Since proteins begin to fold as they are synthesized on the ribosome, two critical events may be recognized. First, folding occurs within the ribosome exit tunnel, a molecular corridor (100Å long, 15Å wide with a 10Å-wide constriction) through which the elongating nascent peptide moves during translation. Second, folding continues as the protein moves through the endoplasmic reticulum (ER) membrane. The ribosome exit tunnel is designed to facilitate nascent protein folding. It has a heterogeneous microenvironment of confined space, non-bulk water^9–11^, and predominantly rRNA. Moreover, the tunnel, has distinctive structural features and ‘constrictions’, including a folding vestibule at the exit port that facilitates peptide folding^12^. Complementarily, the native *protein sequence* of channel domains has evolved to optimize biogenic folding and molecular interactions with the ribosome and ER membrane^12–18^.

**Figure 1.**
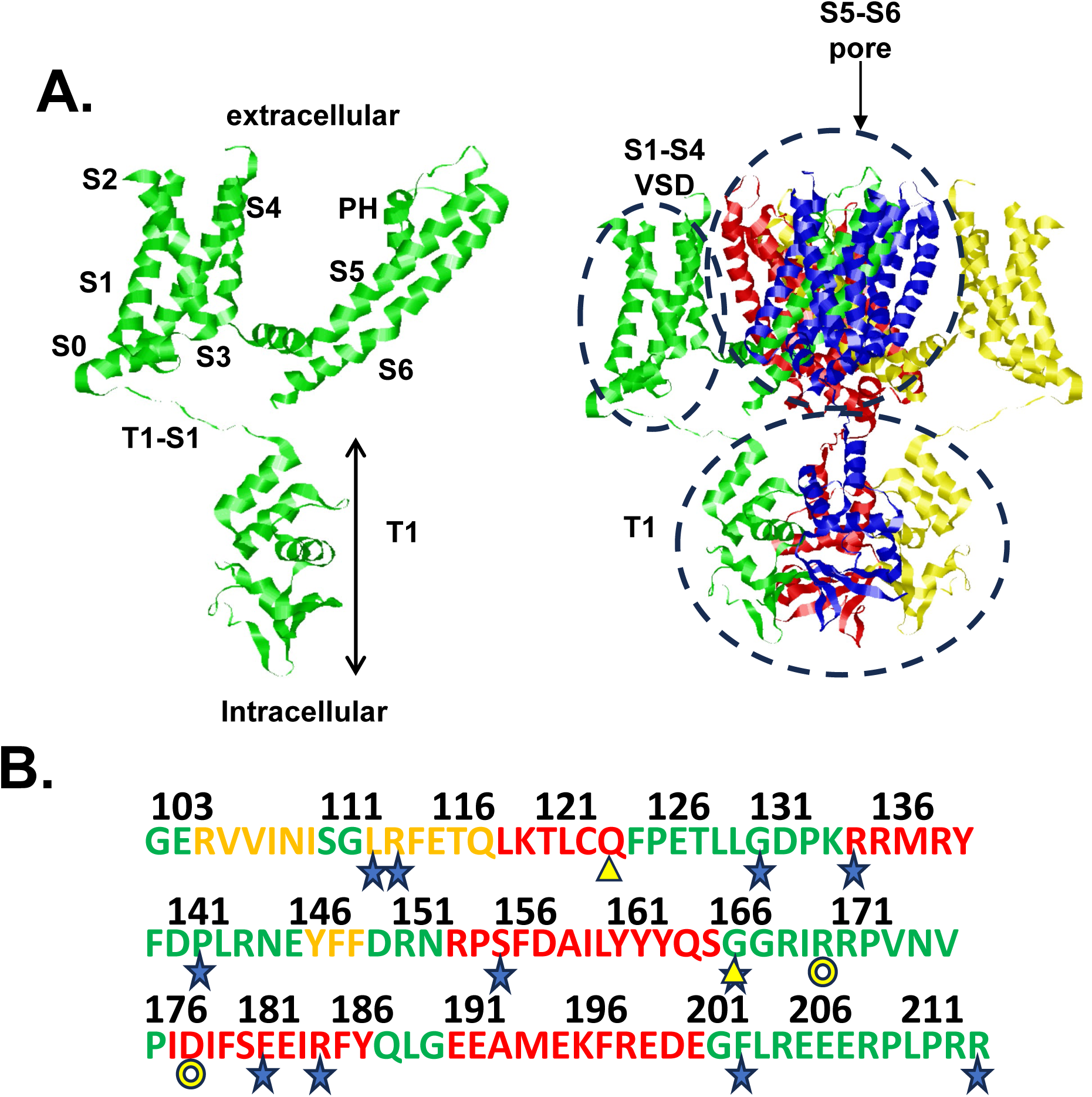
Kv1.3 channel structure and sequence. **A.** Mature Kv1.3 monomer (PDB ID 7SSX^56^ with transmembrane segments, S1-S6, pore helix (PH), cytosolic T1 domain (left) and assembled tetrameric complex (right). Each of the four subunits is shown in a different color. **B.** Location of *KCNA3* variants. Sequence of WT T1. Location of variants are indicated by blue asterisk. Residues R170 and D178, located at T1-T1 interfaces in the correctly folded tetramer, are indicated by yellow circle and were mutated to cysteines for the bifunctional quaternary folding experiments. Residues Q124 and G166, located at an intramolecular T1 tertiary fold, are indicated by a yellow triangle, and were mutated to cysteines for the bifunctional tertiary folding experiments. Assignments for helix (red), β-strand (yellow), and loop (green) are taken from PDB ID 7SSX^56^.

Kv channels are tetrameric, with each monomer containing six transmembrane segments (S1-S6) divided into a voltage-sensing domain (S1-S4) and a pore domain (S5, pore helix, selectivity filter, and S6; Fig. 1A) used for ion permeation. In the cytosolic N-terminus, Kv1 channels contain a highly conserved sequence, the ‘T1’ domain ^19,20^. T1 serves as an intersubunit recognition domain that ensures correct isoform assembly, docks functionally important auxiliary proteins^21–25^, underlies axonal targeting^26^, and contributes to the gating of the mature channel^27–30^. T1 is equally important in early biogenic events. T1 sequences fold as they emerge from the ribosomal tunnel^14,31^, which enables T1 domains to expediently tetramerize at the ER membrane while cotranslation of the downstream nascent chain continues. These two early events, i.e., tertiary and quaternary T1 folding, are coupled^32^, and stabilized by T1 hairpin subdomains (Fig. 1A) that communicate through intervening H-bonds^28,33^.

The aforementioned early-stage folding events depend on the native protein sequence, which evolved to ensure correct architectural conformations during cotranslation and specific residue interactions for correct function of the mature channel. In this study, we focus on a prototypical *Shaker*-like channel, human Kv1.3 (Fig. 1A, encoded by the *KCNA3* gene), which is highly expressed in nerve and immune cells. Kv1.3 is essential for T-lymphocyte activation and proliferation^34–38^, is linked to chronic inflammatory disease, autoimmune and metabolic disorders, attention deficit, and has been implicated in altered insulin sensitivity, neoplastic behavior and malignancy, and modulation of apoptosis^39–41^. The severity of many of these pathologies correlates with Kv1.3 expression at the plasma membrane^39–43^; yet the underlying nascent protein folding/assembly defects that may contribute to altered expression have not been identified. It is thus compelling to ask whether *KCNA3* T1 variants associated with human disease introduce early-stage folding defects.

## RESULTS

If T1 is to serve as a recognition domain for tetramer assembly or a scaffolding function for auxiliary proteins or to direct Kv channels to specific cellular locations, then genetic variants in T1 may undermine the biogenic folding efficiency and consequently, the functional expression of Kv1.3. The genetic basis of human disease has traditionally been investigated using a “phenotype-first” approach, i.e., patients with known phenotypes are gene sequenced to discover potential underlying variants that might be responsible. Instead, we used the recently developed unbiased “genome-first” approach^44–46^ to identify specific *KCNA3* rare variants clustered in the highly structured and conserved T1 domain and associated with clinical consequences. Pilot exome analysis for >40,000 unselected patients in the Penn Medicine Biobank (PMBB, University of Pennsylvania), which are linked to the electronic health record (EHR), identified 20 missense variants, predominantly in the T1 domain, with a high probability of being deleterious (Revel scores 0.5-0.9). Analysis of the combined effect of all 20 variants (burden analysis) revealed association with a wide range of pathological phenotypes in EHR (e.g., Hodgkin’s disease, immune deficiencies, and cachexia).

Focusing on variants in the T1 sequence (Fig. 1B, Supplementary Table 1), we carried out exome analyses of *KCNA3* in the PMBB Database, the UK BioBank TOPMed DataBase, ClinVar DataBase, and the VarSite DataBase. The latter is a predictive software that uses genomic features and structural information from the Protein Data Bank to quantify the probability of a deleterious effect and disease association. The structural location of these variants in the T1 domain of the mature WT sequence are shown in Figure 2, revealing multiple intra- and intersubunit interactions within ≤ 4.5 Å. All these variants have high CADD PHRED and Revel scores (scores >20 and ≥ 0.5, respectively), which are deleterious scores and implicate a disruptive effect on protein folding. Several native residues (e.g., R114, R135, E182, R185) exhibit electrostatic and H-bond interactions with neighboring residues, which contribute to the stability of the folded T1 domain in the native WT. The variants at these positions (R114G, R135L, E182K, R185L) disrupt these interactions. Note also that E116 has both intramolecular and intermolecular interactions with residue R114, an electrostatic interaction that likely is eliminated in the variant R114G.

**Figure 2.**
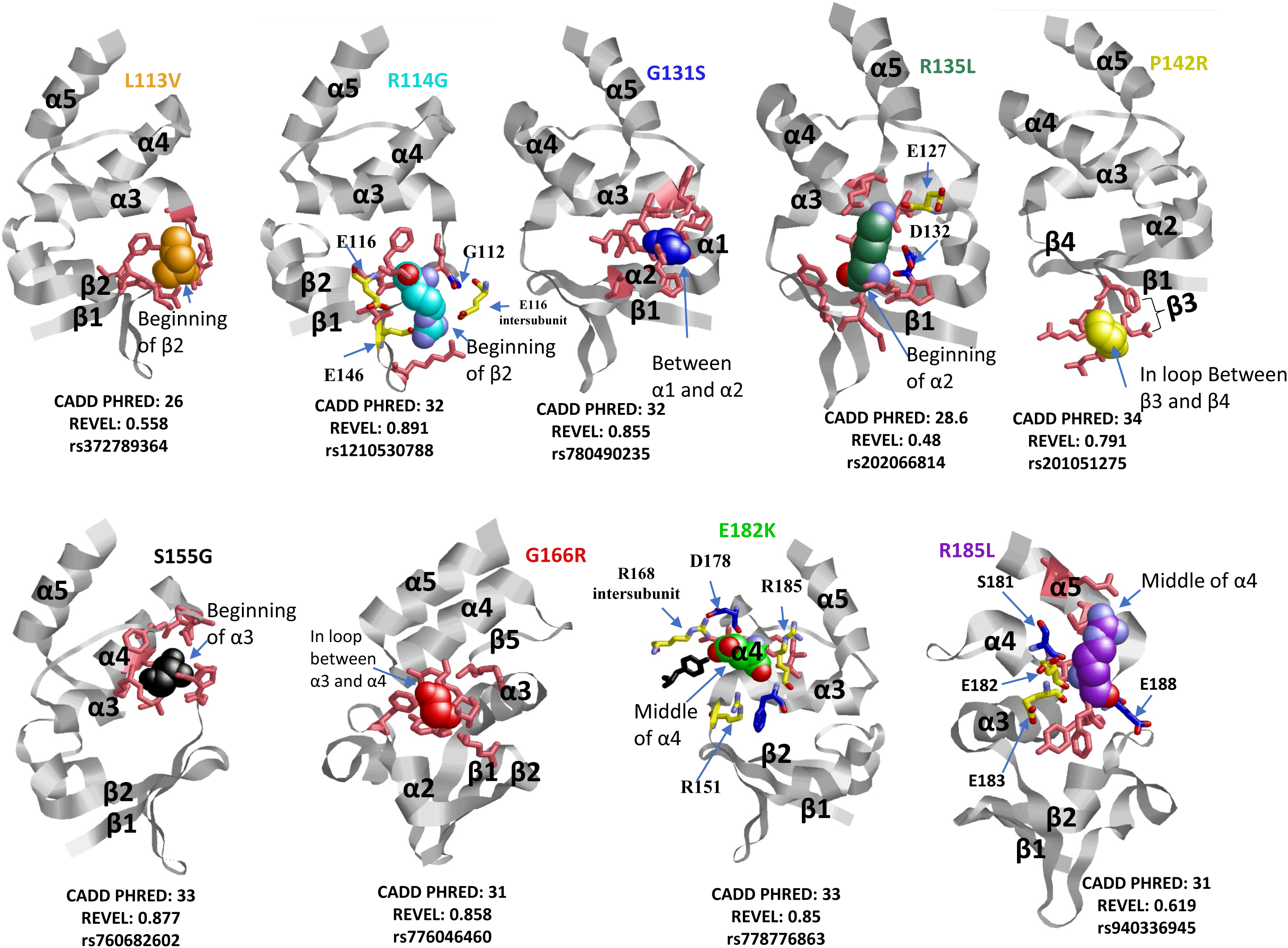
Images of the T1 folded domain. Mature Kv1.2 (PDB ID 1QDV^28^) was used to highlight location of each variant. All amino acid numbers are equivalent residues in Kv1.3. The native residue at the site of the specified variant is depicted as a spacefilled model. Residues within 4.5 Å of the variant are shown as sticks (intra- and intersubunit in pink and black, respectively). Blue sticks represent H-bonds, yellow sticks represent ion-ion interactions. CADD PHRED and Revel scores for each variant are shown beneath each structure and in Supplementary Table 1. Rendered with RasMol.

To investigate the association between these specific genetic variants and a wide array of phenotypes (PheCodes) across genetically inferred populations of European and African ancestry from the Penn Medicine BioBank data, we used the Scalable and Accurate Implementation of Generalized mixed model (SAIGE), a comprehensive analytical approach. Three variants (E182K (rs778776863), F203S (rs199864361), R214C(rs145190352)) were analyzed (rs ID, reference single nucleotide polymorphism, listed in Supplementary Table 1).

Our ancestry-stratified PheWAS analysis identified significant associations between the rare variants and multiple phenotypes across European and African ancestry populations in the Penn Medicine Biobank. These analyses yield a p value and a beta value for each phenotype association. The beta coefficient represents the effect size of the variant on the phenotype, indicating the strength and direction of the association. Positive beta values indicate that the minor allele is associated with increased phenotype measure or risk, while negative values suggest a protective effect. The absolute value of beta reflects the magnitude of the variant’s influence, with larger absolute values indicating stronger effects. Variant R214C in particular showed a strong association with the following phenotypes in the European ancestry cohort (EUR) : diverticulitis (p value = .00019, beta=0.91), mental disorders during/after pregnancy (p value = .0002, beta=0.73), diverticulitis of small intestines (p value =.00025, beta=1.3) congenital anomalies of great vessels (p value =.00044, beta=1.8), and heart block (p value =.00072, beta=1.9) (Fig. 3). In the African ancestry cohort (AFR), R214C was significantly associated with mental disorders after/during pregnancy (p value =.00059, beta=5.0). However, F203S and E182K variants showed modest associations (p values of .006 and .001, respectively; See Supplementary Figure S1 for specific clinical phenotypes; see Supplementary Figs. S2, S3, S4 for an overview of all tested phenotype categories). Within the AFR ancestry only, F203S has significant associations with numerous phenotypes (Supplementary Fig. S1A). The most notable association was with congenital disease, specifically valvular heart disease and heart chamber anomalies (p value=.006, beta=4.9). This association was not observed in non-AFR populations, highlighting the genetic variant’s particular relevance to the AFR group.

**Figure 3.**
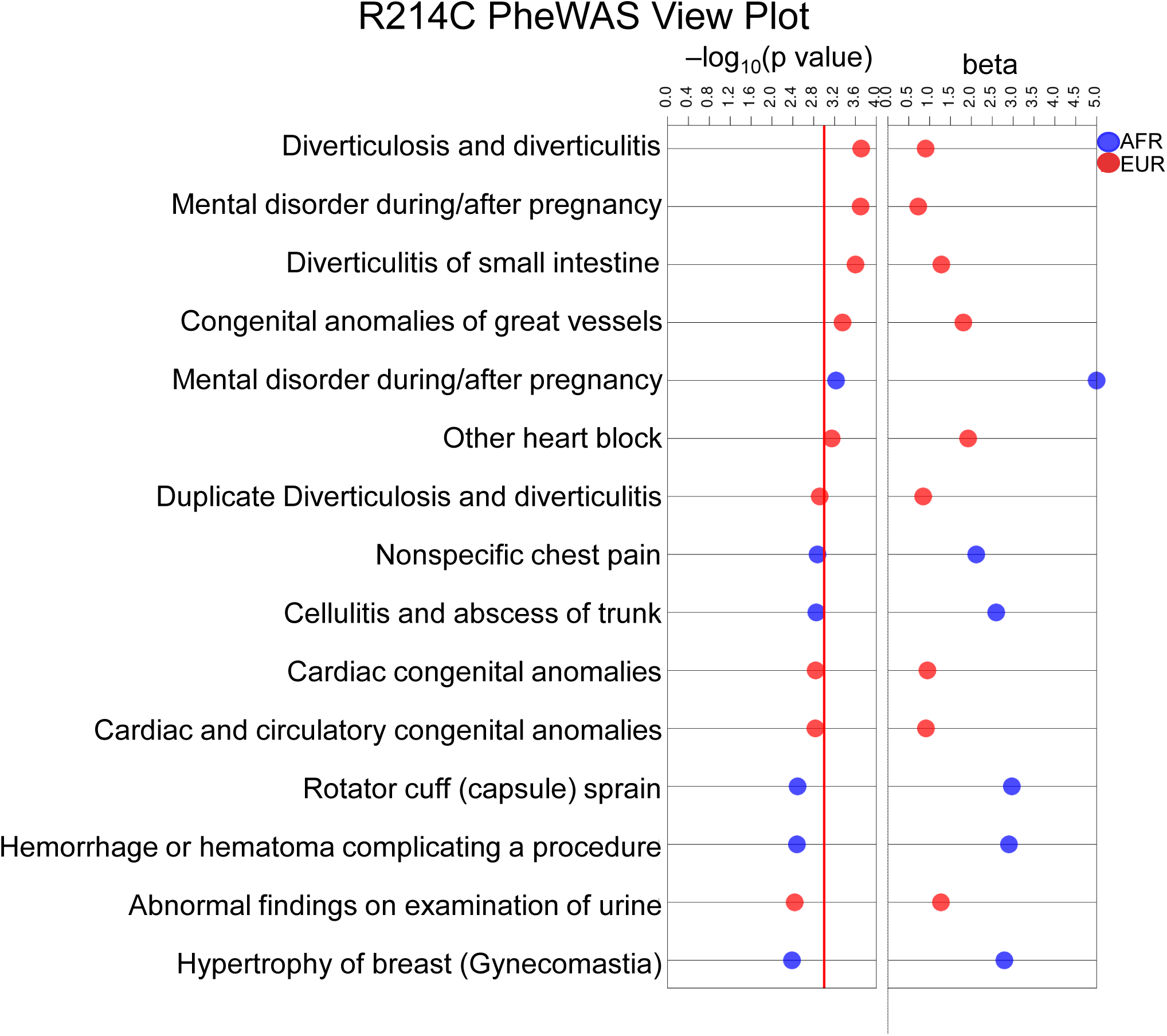
PheWAS-view plot for variant R214C. Data analysis shows −log_10_ p value (left) and effect size estimates (beta) from PheWAS analyses on the x-axis and specific clinical phenotypes on the y-axis. This plot shows top 15 results from PheWAS analyses. Red line shows the p value=.001. AFR results are shown in blue points and EUR are shown in red points. For broad phenotype categories, see Supplementary Figures S2-S4.

The observed phenotype associations and deleterious predictions of all 11 variants prompted us to probe differences in folding and assembly efficiencies. First, we determined the probability of nascent proteins to associate with the microsomal membranes and be glycosylated. Second, we assessed the probability of intermolecular tetramer formation and intramolecular tertiary folding of T1. As shown in Figure 4A, the efficiency of membrane insertion is decreased for variant R114G (0.42±0.02 (n=5)) compared to native WT (0.57±0.03 (n=15)), with a concomitant decrease in fractional N-linked glycosylation (R114G, 0.21±0.02 (n=5) versus WT, 0.34±0.02 (n=23)). The range of fractional glycosylation, which is typically dependent on the relative quality of the microsomal membranes, the distance between the glycosylation site and the intraluminal membrane, and the efficiency of the oligosaccharyltransferase, is 20-40%. Nonetheless, N-linked glycosylation is an indicator of protein insertion into the microsomal membrane. These findings indicate that expression of variant R114G yields ∼ 50% glycosylated membrane-associated Kv1.3 compared to WT.

**Figure 4.**
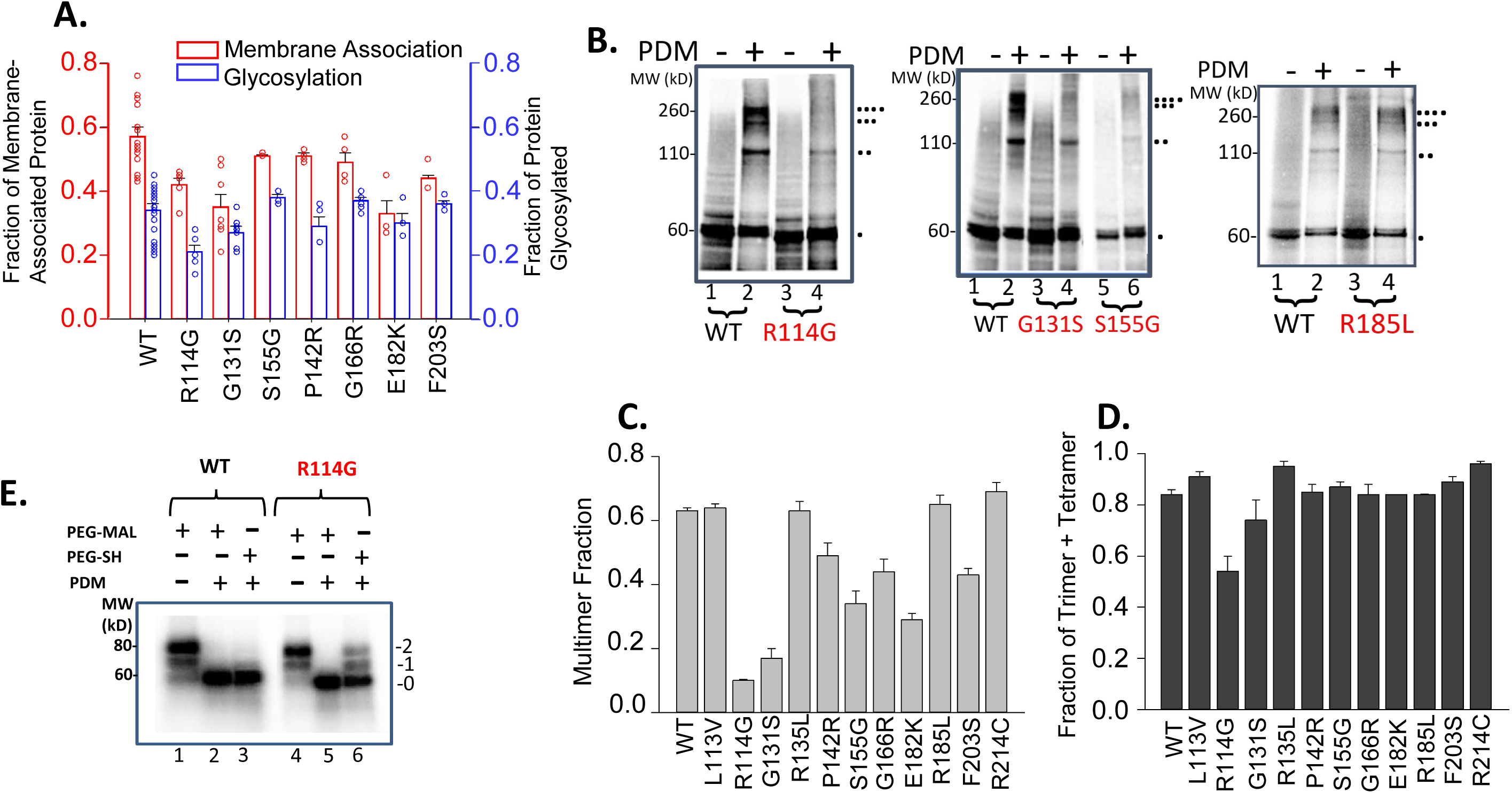
Kv1.3 membrane association, core glycosylation, and T1 folding. **A.** Membrane-Associated Kv1.3 and core glycosylation. Membrane association efficiency of full-length Kv1.3 protein (red bars, left axis) for the indicated variants after 120 min translation reactions in the presence of microsomal membranes. Fraction core glycosylated is depicted as blue bars (right axis) for each variant. Dots are data points; colored bars are means; error bars are ± SEM. **B.** Quaternary folding of T1 domains. Multimer formation for WT (lanes 1, 2) and *KCNA3* variants (red labels, lanes 3-6). Modification with o-phenyldimaleimide crosslinker (PDM), yields monomer, dimer, trimer, and tetramer, indicated by the number of asterisks, respectively. Two cysteines in the T1 –T1 interface were engineered for this purpose (Fig.1B). **C.** Quantitative analysis of crosslinking gels. Each bar is the fraction of total protein crosslinked as dimer, trimer, tetramer divided by the sum of monomer, dimer, trimer, and tetramer (n=3, mean ± SEM), i.e., equivalent to the probability of crosslinking (*Pxlink*). **D.** Trimer + Tetramer fraction of crosslinked protein. This fraction is calculated as the sum of trimers+ tetramers divided by the sum of dimers, trimers, and tetramers. **E.** Tertiary folding of T1 domains. Tertiary folding of the T1 WT (lanes 1-3) and R114G variant (lanes 4-6). Full-length constructs containing an engineered cysteine pair (Fig. 1B) were translated, then treated with bismaleimide PDM (0.5 mM, Kosolapov and Deutsch, 2003^13^) and finally pegylated with PEG-MAL or PEG-SH (5 mM, 1 hour at 4°C). Numbers to the right represent doubly (2) or singly (1) pegylated or unpegylated (0) protein. *Pfold* is 0.90 for WT, 0.55 for R114G.

To assess the probability of quaternary folding of T1 domains (*Pxlink*) in Kv1.3 nascent proteins, we determined the multimerization efficiency of Kv1.3 nascent subunits for each variant. Engineered cysteines (170 and 178) in adjacent subunits at the subunit-subunit interface can be crosslinked with ortho-phenyldimaleimide (o-PDM)^31,32^(Fig.4B). *Pxlink* is the fraction of total protein that is crosslinked to give dimers, trimers, and tetramers. It can be calculated from quantitation of the corresponding bands on a protein gel, yielding a *Pxlink* of ∼0.60 for WT T1. *Pxlink* for variants is shown in Figure 4C. Variants R114G, G131S, P142R, S155G, G166R, E182K, and F203S, manifest decreased quaternary folding of T1 domains. Additionally, several of these variants may induce electrostatic perturbations in the folded T1 domain thus leading to inefficient tetramerization. Among the variants, R114G, G131S, E182K, and F203S each impair tetramerization and membrane association (Fig. 4A,C). Moreover, the fraction of crosslinked protein that is captured as trimers and tetramers, i.e., (trimers+ tetramers)/(dimers + trimers + tetramers) (Fig. 4D) reflects the efficiency of stepwise multimerization. A ratio close to 1 indicates that dimer concentration is minimal due to efficient dimerization of dimers, the main mechanism by which Kv1.3 tetramers form^47^. R114G has a ratio ∼ 0.5, suggesting a reduced efficiency of higher order multimerization with an accumulation of crosslinked dimers. Figure 4D suggests that variant R114G has a lower probability of forming trimers and tetramers, and generates a more stable dimer, perhaps consistent with tetramer instability observed in MD simulations (below, Figs. 5, 7).

**Figure 5.**
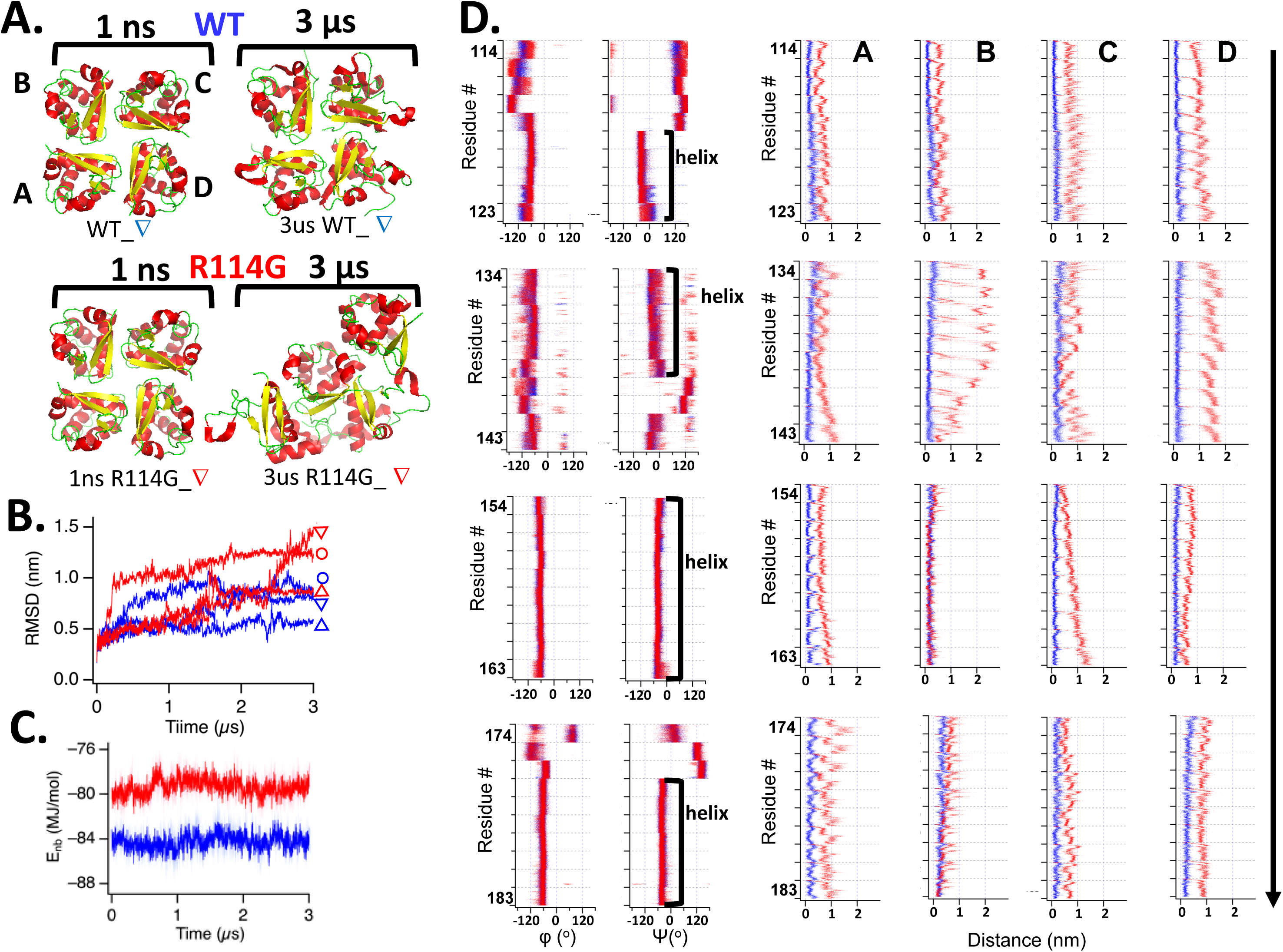
Structural and energetic features of WT and R114G T1 tetramers. **A.** Tetrameric T1 structures derived from MD simulations for WT and R114G at 1 ns and 3.0 μs viewed from the intracellular side. In each subunit (A, B, C, D), helices are colored red, β strands yellow, loops green. Structures are rendered with PyMol. Symbols (D, Δ) represent one of three independent MD simulations of WT and R114G, respectively (See Supplementary Fig. S5A for all three simulation-based structures for WT and for R114G). **B.** Root-mean-square deviations (RMSDs) of non-hydrogen atoms from three independent WT (*blue*) and three independent R114G (*red*) simulations. Symbols to the right of the traces indicate different simulation runs. **C**. Mean nonbonded potential energies (E_nb_) of the WT (*blue*) and R114G (*red*) T1 tetramers from three independent simulations (n=3 for each curve) was calculated as the sum of the van der Waals and electrostatic energies. Lightly shaded areas represent standard deviation. **D.** Dihedral angles and distance moved for WT and R114G variants. *Left*: Changes in φ and ψ angles for select residues in a subunit from one illustrative WT system (*blue*) and a subunit from one illustrative R114G system (*red*) during the 3-µs simulation are shown superimposed. For each residue indicated along the ordinate, time progresses from 0 to 3 µs downward. Secondary conformations are assigned according to PyMol as in Fig.1B. *Right*: Distance moved for the α-carbon of each selected residue from its initial location at 1 ns to its location at 3 μs in the four subunits (A, B, C, D) in the systems shown in (**A**). Vertical arrow indicates that each plot progresses from N-terminal to C-terminal residues. The measured intersubunit distance for crosslinkable cysteines at R170 and D178 was 7-10 Å for WT vs. 25-27 Å for variant R114G.

Quaternary folding is intimately linked to secondary and tertiary folding of the protein, as we have previously shown for folding and function of the T1 domain^32^. To estimate the probability of tertiary folding of T1 (*Pfold*), we used PDM crosslinking and pegylation mass-tagging to calculate a probability of *<u>intra</u>*molecular tertiary folding^14–16^. This assay (Fig. 4E) completely obviates capturing rare conformations, a liability of disulfide-capture strategies. Both *folded* and *<u>un</u>folded* species are measured simultaneously due to PDM’s irreversible and rapid crosslinking of Cys pairs at a folded peptide interface (1 PDM/Cys pair) and covalent modification of the individual cysteines in an unfolded peptide (1 PDM/Cys). Guided by X-ray structures (aKv1.1^33^) and function, we engineered Cys pairs at tertiary intramolecular interfaces (residues Q124C/G166C). Defective tertiary T1 folding occurs in the full-length, membrane-associated variant R114G compared to WT (Fig. 4E): WT = 0.89±0.01, R114G = 0.59±0.04, n=3. This variant defect is already manifest in the nascent peptide attached to the ribosome derived from a BstEII-cut mRNA construct (*Pfold:* WT = 0.67±0.01, R114G = 0.38±0.02, n=2).

To understand the energetic basis for variant folding defects, we performed all-atom molecular dynamics simulations of the intracellular T1 tetramer of WT and variant R114G. To facilitate comparison of the computational studies and the experimental results presented above on time scales of minutes to hours, the simulations were performed at 90 °C. Such high temperature simulations may better reveal the structural stability differences between the WT and R114G complexes. Figure 5A (and Supplementary Figure S5A) shows the tetrameric T1 structures of WT and R114G variant at 1 ns and a final point of 3 μs. The WT tetramer undergoes subtle changes in loop and helical arrangements whereas the R114G undergoes much more dramatic changes of secondary and tertiary structures in each of the four subunits. While subunit asymmetry is apparent in both WT and variant dynamics and conformations, the R114G T1 tetramer is clearly more unstable than the WT T1 tetramer and shows marked unfolding of each subunit and the quaternary conformation of the tetramer.

The structural stability differences between WT and R114G are also evident in the root-mean-square deviation (RMSD) plot (Figure 5B), which shows the RMSD of non-hydrogen protein atoms compared with the respective starting tetrameric structures. Two of the three R114G complexes (Figure 5B, *red*) exhibited noticeably greater increases in RMSD compared with the WT (Figure 5B, *blue*) by the end of the simulation epochs, indicative of diminished structural stability of the T1 complex by the mutation R114G. Consistent with the geometrical differences noted, the nonbonded potential energies, comprised of van der Waals and electrostatic interactions, of the R114G tetrameric complexes studied were noticeably greater than those of the WT complexes (Figure 5C). When the nonbonded energies of the individual subunits were compared, similar differences were also evident (Supplementary Fig. S5B), with R114G subunits having higher energy levels.

The macroscopic geometrical and energetic stability differences between WT and R114G should be reflected in the secondary conformations, in part characterized by dihedral angles. Variability in the protein backbone dihedral angles is expected to serve as an estimator of secondary structure stability; greater variability suggests less stable secondary structures. The angle variability in WT and R114G was quantified using the standard deviation (SD) of dihedral ɸ (N-C_α_) and ψ (C_α_-C) values during the simulation (Fig. 6). An SD value was calculated for each subunit, with 12 subunits analyzed for WT (4 subunits in 3 complexes) and 12 for R114G. The SD results are strikingly informative. First, while the SD values of some helical segments (e.g., 152-162, 175-185) are very small and virtually indistinguishable between WT and R114G, those for helical residues in 120-125 and 135-142 are distributed over an identical but very wide range for both WT and R114G. Second, residues in the region of the variant R114G (residues 110-116) show large fluctuations in both ɸ and ψ angles with SDs averaging ∼30° and 100°, respectively whereas the WT residues in this interval are all uniformly and tightly clustered with SDs of ∼15° for both ɸ and ψ. Third, subunit asymmetry in dihedral angle variation within the tetramer is apparent, e.g., 4 identical symbols for a given residue in each subunit of a single tetramer are unequally distributed over a range of dihedral angles. Fourth, the final helical segment at the C-terminus of T1 (residues 191-200) is more mobile than is typical for an α-helix due to its being untethered from the T1-S1 linker and the rest of the channel (Fig. 1A).

**Figure 6.**
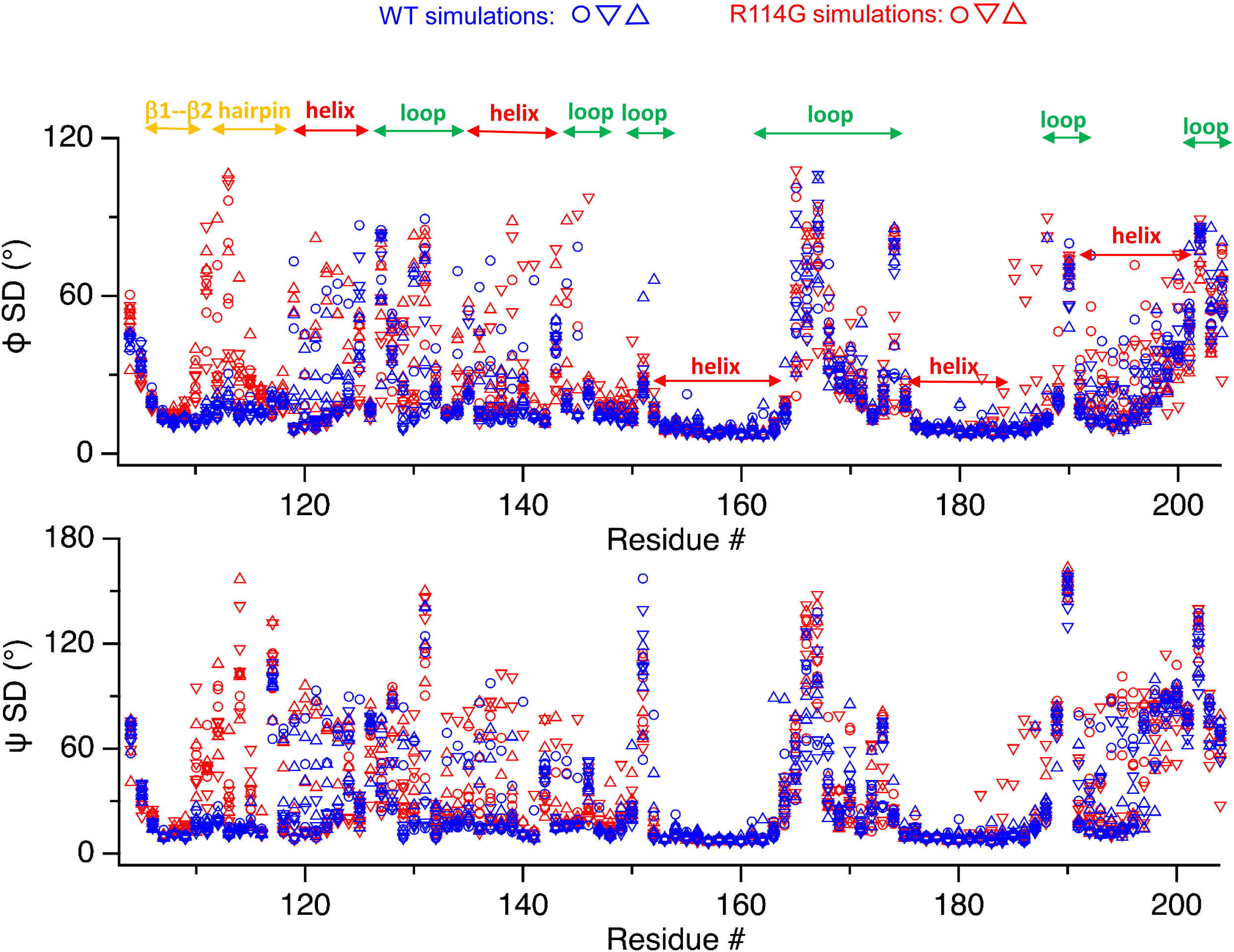
Variability in dihedral angles for WT and variant R114G. Variability (SD) in phi/psi angles (φ, ψ) of residues for each subunit within the 3-μs simulation in three independent MD simulations of WT 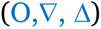 and R114G 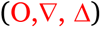. Each complex has 4 subunits and thus there are four symbols of one type and one color at each residue.

We next addressed how the secondary structures characterized by ɸ and ψ angles relate to the overall tertiary structural instability. Figure 5D compares the ɸ and ψ angles derived from MD simulations for select residues in helical segments (left) with their C_α_ displacements from the initial positions at 1 ns (right) over 3 μs for WT (*blue*) and R114G (*red*). The dihedral angles for WT and R114G in these helical segments remain similar, indicating that the helical secondary structures are intact in both WT and R114G throughout the simulation duration. In sharp contrast, the R114G C_α_ displacements from their initial positions (right, *red*) are markedly greater than those of WT (right, *blue*); a large fraction of the secondary structure survives the disruption of the native tertiary interactions, i.e., tertiary structures unfold first, which is readily seen in the simulated R114G structures depicted in Figure 5A. The negligible displacements of the helical C_α_ atoms from the initial positions in the WT T1 subunits also suggest that the initial four-fold symmetry of the WT tetrameric T1 complex is well maintained. Consistent with these results, structures derived from MD simulations for 1-3000 ns (Fig. 5A, Supplementary Fig. S5A) in R114G manifest the induced consequences of these dramatic changes in tertiary folds and the α-carbon distances moved. Another intriguing observation is again the asymmetric temporal and spatial dynamics of the subunits (labeled A, B, C, D), which is most striking for the variant (Fig. 5D, right, *red*). The role of subunit asymmetry in biogenic assembly of oligomeric complexes is unknown and warrants further investigation.

The simulated WT T1 structure suggests that the N-terminal β-strand segment encompassing residues 111-117 may be important for intersubunit interactions and structural symmetry. Thus, the nonbonded potential energy of these residues was calculated for each subunit in WT and R114G (Fig. 7A). In R114G complexes, these residues possess greater nonbonded energy than in WT, illustrating the destabilizing influence of the mutation on this segment and the overall structure.

**Figure 7.**
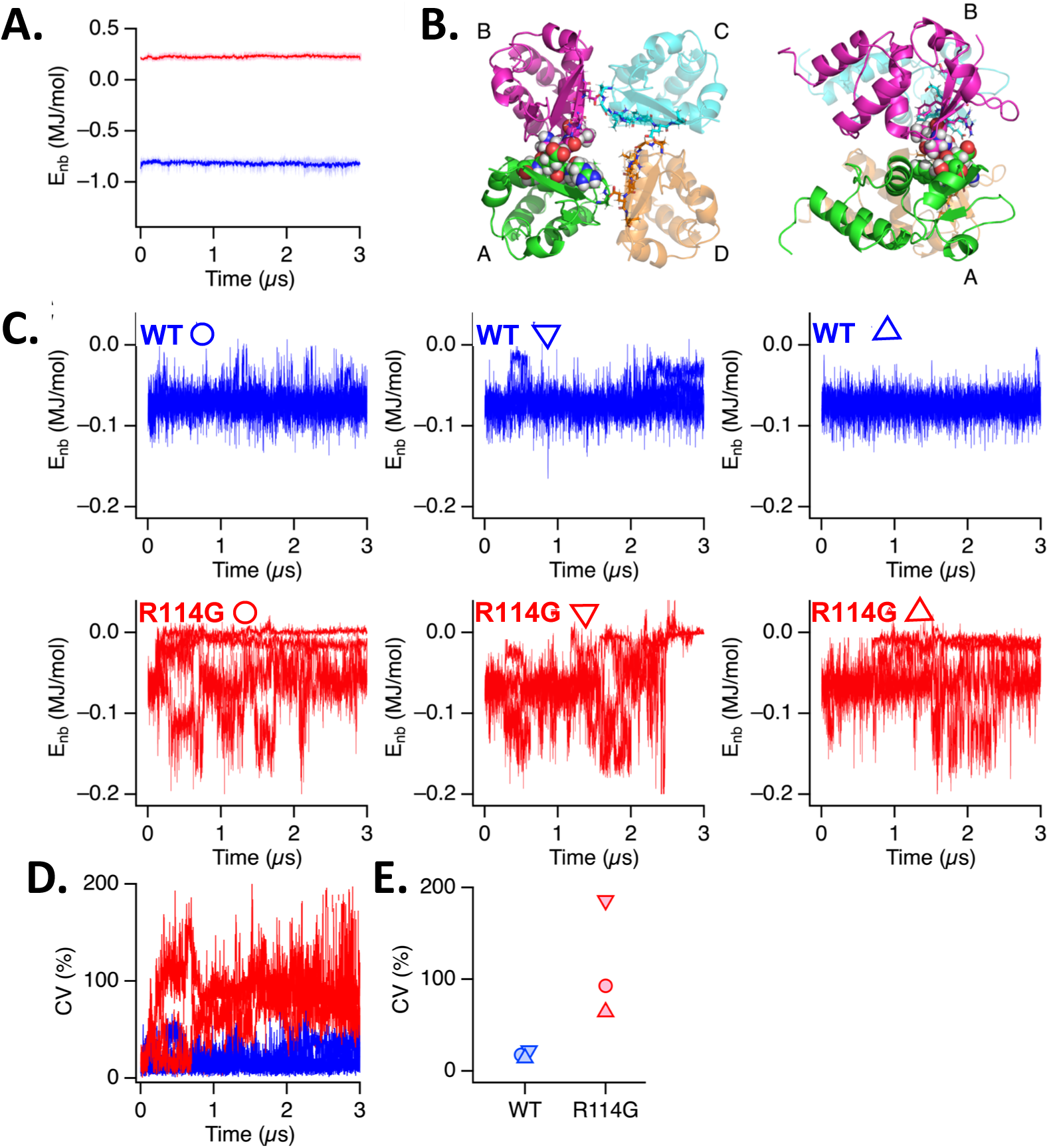
Intersubunit interactions mediated by the N-terminal residues. **A**. Mean nonbonded energy (E_nb_) levels of the residues 111 through 117 in WT (*blue*) and R114G (*red*) across the four subunits during three simulations. Thus, each curve represents the mean of 12 (E_nb_) at each time point. Light shaded areas represent standard deviation. **B**. T1 domain (PDB 7SSX) view from the intracellular side (*left*). Subunits are rendered in different colors with residues 114–117 in subunit A (*green*) and 111–113 in subunit B (*magenta*) shown using spheres. The same residues in the remaining subunits are depicted as sticks. On the right, the structure on the left is rotated 90°along the Y-axis so that the membrane is on the left and the intracellular side is on the right of the image. **C**. Nonbonded interaction energies between residues 114–117 in one subunit and residues 111–113 in the adjacent subunit (clockwise, viewed from the intracellular side) as a function of simulation time. Each graph represents the four interaction energies within a tetrameric complex from a separate simulation run (WT: blue, R114G: red). **D**. Coefficient of variation (CV) of the intersubunit interaction energies from the three WT (*blue*) and three R114G (*red*) simulations shown in C. Some extreme values >200% in one R114G simulation are not shown. **E**. Time-averaged CVs for the WT (*blue*) and R114G (*red*) simulations. Symbols (O,∇, Δ) used in C. and E. correspond to those used in Figures 5A, 5B, 6, and Supplementary S5A to indicate different simulation runs.

Within this β-strand, residues R114 through T117 in one subunit make close contacts with residues S111-L113 in an adjacent subunit in WT (clockwise viewed from the intracellular side; Fig. 7B). R114 may also contribute to intrasubunit interactions. In WT, the subunit interaction energies among the four subunits in each system were nearly identical and remained stable, showing only small variability, throughout the simulation (Fig. 7C). In contrast, the subunit interaction energies in R114G were far more variable, both among the four subunits and with time. The interaction energy variability among the four subunits in each simulation was characterized by the time-averaged coefficient of variation (CV, Fig. 7D). Every R114G system had a much greater CV value than any of the three WT systems (Fig. 7E). Thus, the unmatched and asymmetrical interaction energies may be associated with the greater structural instability observed with R114G.

## DISCUSSION

Nascent proteins fold during all stages of biogenesis. This process is particularly complex for ion channels composed of multiple distinctive biogenic and functional domains and subunits^12–16,24,31,32,48^. For Kv1.3, specific T1 residues are critical for tertiary and quaternary structure formation during biogenesis ^28,32^, which are highly coupled processes with a Pearson correlation coefficient of 0.86^32^. The monomeric T1 domain cannot oligomerize in the absence of a threshold level of tertiary structure. Given these molecular determinants, we set out to explore whether human native *KCNA3* variants with predicted deleterious consequences and clinical phenotypes might manifest folding defects early in the biogenesis of the nascent peptide. Our folding assays indicate markedly different folding probabilities for several of the native variants compared to WT, providing strong evidence for our hypothesis: native *KCNA3* variants in the T1 domain impair early-stage quaternary and tertiary Kv folding and membrane insertion. Electrostatic, H-bond, and van der Waals interactions at the T1-T1 interface and/or within a T1 domain may energetically contribute to tetramer formation and are therefore compromised in selected variants. Those variants that exhibit decreased *Pxlink* are consistent with the deleterious predictions shown in Supplementary Table 1. However, one variant predicted to be functionally deleterious yields a similar *Pxlink* to WT (Fig. 4B), which warrants further investigation.

Despite the well-known limitations of MD simulations, including challenges in fully describing electrostatic interactions and in relating short timescale simulations to biochemical experiments occurring on longer timescales, our simulation results do provide multiple insights that may bring us closer to understanding the molecular mechanisms underlying inefficient folding. The R114G variant produces a readily observable, more unstable T1 domain; changes in its quaternary and tertiary structures are obvious in at least two of the three simulation runs (Fig. 5B). While some helical segments remain relatively stable in both WT and R114G (Fig. 5D, *left*), the R114G variant increases fluctuations in atomic positions at select residues, including near position 114 (Fig. 6). Considering the aforementioned pre-requisite for a threshold amount of tertiary structure for proper oligomeric formation^32^, the energetic instability of the variant probably impairs formation of a symmetrical tetrameric complex and promotes tertiary unwinding (Fig. 5D, *right*). Moreover, unfolding of each subunit within the variant tetramer is not identical. In addition to tertiary level impairment, perhaps the most striking finding is the non-equal asymmetrical interactions among the four subunits caused by the variant R114G. In WT, the four intersubunit interactions mediated by residues 111-113 and 114-117 are nearly equal and stable over time. In contrast, in R114G, the interactions vary widely and fluctuate noticeably over time (Fig. 7C, D), distorting the symmetry and stability of the quaternary structure.

Our folding experiments and molecular simulations were initially motivated by genetic analyses, which revealed associations of variants with numerous clinical phenotypes. Several intriguing propositions arise from these observations of a variant’s rarity and the gene’s intolerance to missense variations. The numerous clinical phenotypes also imply pleiotropy. For example, R214C in the EUR population, has a minor allele frequency (MAF) of 0.0019, corresponding to approximately 1 in 263 individuals carrying the minor allele. The variant is even rarer in the AFR population, with a MAF of 0.00044, or about 1 in 1,136 individuals. The functional importance of this variant is further supported by its missense score of 6.43 (Z-score) based on Gnomad data. A Gnomad Z-score is a measure of genetic constraint, indicating how tolerant a gene is to missense variations. This high positive Z-score indicates that the gene is under strong selective pressure to maintain its native protein sequence, with fewer missense variants observed than expected under neutral evolution. Such a score suggests that missense changes in this gene are likely to have significant functional impacts on the encoded protein. This evidence of genetic constraint reinforces the potential biological significance of *KCNA3* variants and may explain its associations with multiple phenotypes. The combination of the variant’s rarity and the gene’s intolerance to missense variations underscores the importance of further investigation into the specific molecular mechanisms by which a variant might influence the observed phenotypes.

Among the clinical phenotypes significantly correlated with E182K, F203S, and R214C, are broad categories of congenital disease, digestive disease, neoplasm, nerve disease and pain, and respiratory disease. This finding suggests a potential pleiotropic effect, possibly reflecting redundancy or compensatory mechanisms induced by a disruptive mutation and merits further investigation. The SAIGE analyses of other genetic variants were too low in frequencies to be effectively tested within the scope of PheWAS. This may reflect a potential evolutionary significance whereby exceedingly rare variants confer a significant survival disadvantage or are otherwise deleterious, thereby being rapidly removed from the gene pool through natural selection processes. This observation hints at the intricate balance between *KCNA3* genetic variation and its impact on human health. Understanding how these rare variants influence diverse phenotypes could provide valuable insights into the shared pathological pathways, potentially opening new avenues for therapeutic interventions.

## METHODS

### Constructs and *in vitro* translation

Standard methods of bacterial transformation, plasmid DNA preparation, and restriction enzyme analysis were used. The nucleotide sequences of all mutants were confirmed by automated cycle sequencing performed by the DNA Sequencing Facility at the University of Pennsylvania School of Medicine on an ABI 3730XL Sequencer using BigDye terminator chemistry (A0BI). Engineered cysteines and restriction enzyme sites were introduced into pSP/Kv1.3/cysteine-free using QuikChange Site-Directed Mutagenesis Kit (Agilent, Santa Clara, CA). All mutant DNAs were sequenced throughout the region of the mutation. Capped cRNA was synthesized *in vitro* from linearized templates using SP6 RNA polymerase (Promega, Madison, WI). Linearized templates for Kv1.3 translocation intermediates were generated using several restriction enzymes to produce different length DNA constructs lacking a stop codon and to position the nascent peptide segments at the specified locations. Proteins were translated *in vitro* with [^35^S]methionine (2 μl/25 μl translation mixture; ∼10 μCi/μl Express (EasyTag), Revvity Health, Boston, MA) for 1 hr at 22°C or 2 hr at 30℃ in a rabbit reticulocyte lysate (Promega, Madison WI; 2 mM final [DTT]) with 1 μl of microsomal membranes (canine pancreas: gift from Gunnar von Heijne; Promega, Madison WI) when applicable according to the Promega Protocol and Application Guide.

### Genetic exome analysis

We investigated exome analysis for 45,000 unselected patients from a recently expanded Penn Medicine Biobank (PMBB) database. The PMBB screen yielded 20 new Kv1.3 candidates. We then screened these 20 candidates across three databases. The first was the UK BioBank TOPMed DataBase, which reports EHR-derived PheWAS data in 400,000 white British individuals. The second database was ClinVar, which reports clinical significance information for human Kv1.3 variants. The third database was VarSite, a predictive database that uses structural and genomic features from the Protein Data Bank to quantify the probability of a deleterious effect and disease association. The UCSC Genome Browser (genome.ucsc.edu) was used to obtain Revel scores and the VarSite database (ebi.ac.uk/research/thornton/software/) was used to obtain CADD PHRED scores. Pathogenic interpretation of these scores was determined by Ensembl (useast.ensembl.org) and published literature^49,50^. These evaluations prioritized 11 of the 20 PMBB candidates for experimental investigation. The 11 variants are distributed in the T1 domain (L113V, R114G, G131S, R135L, P142R, S155G, G166R, E182K, R185L) and the T1-S1 linker (F203S, R214C). These variants are of interest because they have high CADD PHRED and Revel scores (scores >20 and ≥ 0.5, respectively, are deleterious) and implicate a disruptive effect on protein folding.

### Phenome-wide association analyses

We performed an ancestry-stratified single variant association test for variants of interest using data from the PMBB. The analysis focused on individuals with genetically inferred European ancestry (EUR) and African ancestry (AFR), where principal components derived from principal component analyses were projected on the 1000 Genomes dataset to determine genetically informed ancestry^51,52^. Phenotypes were defined using Phecodes^53^ derived from electronic health records, representing binary health outcomes. To ensure statistical power, phenotypes with fewer than 50 cases were excluded from the analysis, resulting in 1767 Phecodes for testing. Association testing was conducted using the Scalable and Accurate Implementation of Generalized mixed model (SAIGE^54^), which accounts for sample relatedness and can handle case-control imbalance common in large scale genomic studies. The SAIGE model included fixed effects for the genotype of interest and covariates (age at, sex and first five principal components), as well as random effects to account for genetic relatedness. Results below a p value threshold of .01 were considered significant. This comprehensive approach allowed us to identify significant associations between genetic variants and a wide range of phenotypes while accounting for population structure and relatedness within the PMBB cohort. Following the association analysis, we utilized PheWAS view^55^ plots to visualize the results for any significant associations, providing a detailed overview of the genetic variants’ significant associations with top 15 phenotypes.

### Intermolecular crosslinking assay (*Pxlink)*

Translation reactions (25 µl) were carried out in the presence of canine microsomal membranes (1 μl) and rabbit reticulocyte lysate, 2 mM DTT (Promega Protocol and Application Guide) using full-length cysteine-free WT and variant constructs in the background of R170C/D178C constructs. A 7 µl sample was diluted into 700µl of 1x DPBS (Dulbeccos’s, without Ca^2+^ or Mg^2+^, pH 7.3), followed by incubation with 0.5 mM ortho-phenyldimaleimide (o-PDM) for 1 hr on ice. The reaction was terminated by adding 5 mM DTT and incubated at room temperature for 15 min. Then, the reaction was centrifuged through a sucrose cushion (120µl; 0.5 M sucrose, 100 mM KCl, 5 mM MgCl_2_, 50 mM HEPES (pH 7.3) and 1mM DTT) at 4°C for 7 min at 50,000 rpm using a TLA 100.3 Beckman Optima TLX Ultracentrifuge, to isolate membrane-bound peptide. The supernatant was aspirated and the pellet resuspended in 14.5 μl DPBS (1x), 2.5 μl of DTT (10x) and 7 μl of LDS (4x).

### Intramolecular crosslinking assay (*Pfold*)

Translation reactions (25 µl) were carried out in the presence of canine microsomal membranes (1 μl) and rabbit reticulocyte lysate, 2 mM DTT (Promega Protocol and Application Guide) using full-length cysteine-free WT and variant R114G constructs in the background of Q124C/G166C constructs. A 10-20 ul sample of the initial 25-ul translation reaction was added to 500 ul of 1x DPBS buffer containing final concentrations of 4 mM MgCl2 and 2 mM DTT. The suspension was centrifuged at 70,000 rpm for 20 mins at 4°C or 50,000 rpm for 7 min at 4°C through a sucrose cushion (120µl; 0.5 M sucrose, 100 mM KCl, 5 mM MgCl_2_, 50 mM HEPES (pH 7.3) and 1 mM DTT) with a TLA 100.3 Beckman ultra-centrifuge rotor at 4 °C to isolate ribosome-bound peptide or membrane-bound peptide, respectively. The supernatant was removed and the pellet was resuspended on ice in 500 μl DPBS followed by incubation with 0.5 mM o-PDM for 30 min on ice. The reactions used for subsequent pegylation with PEG-MAL were quenched with 10 mM β-mercaptoethanol at room temperature for 10 min, while reactions used for subsequent PEG-SH labeling were not quenched. Samples were layered on 120 ul sucrose cushion, centrifuged at 70,000 rpm at 4°C for 20 min or 50,000 rpm at 4°C for 7 min, resuspended in 50 μl DPBS containing 1% SDS and incubated at room temperature for 30 min. All SDS-treated samples were diluted with 50 μl DPBS buffer containing methoxy-polyethylene glycol maleimide (PEG-MAL, MW 5000; SunBio Korea) for a final concentration of 5 mM or 50 μl of DPBS buffer containing methoxy-polyethylene-thiol (PEG-SH, MW 5000; SunBio Korea) for a final concentration of 5 mM, and incubated on ice for 2 hrs. After pegylation, samples were precipitated with acetone containing 0.4 mM HCl overnight. Pellets were dissolved in 14.5 μl DPBS buffer, 2.5 μl of DTT (10x), and 7 μl of LDS (4x). Samples with truncated constructs were treated with 50 ug/ml RNase and incubated at room temperature for 20 min. *Pfold* was calculated as described in Robinson and Deutsch, 2005^32^.

### Membrane association of nascent protein

A 10-µl aliquot from a 25 µl translation reaction containing microsomal membranes (1 μl) was diluted with 170 µl of DPBS buffer, and was then centrifuged through a sucrose cushion (120µl; 0.5 M sucrose, 100 mM KCl, 5 mM MgCl_2_, 50 mM HEPES (pH 7.3) and 1 mM DTT) at 4°C for 7 min at 50,000 rpm using a TLA 100.3 Beckman Optima TLX Ultracentrifuge, to isolate membrane-bound peptide. The supernatant was decanted to another tube, and the pellet was resuspended in 14.5 μl DPBS buffer, 2.5 μl of DTT (10x), and 7 μl of LDS (4x). A third of the decanted supernatant was transferred to another tube, then precipitated with 900 μl of acetone buffer that contained 0.4 mM HCl and stored at –80°C overnight. After overnight storage, the supernatant acetone mixture was spun down in an Eppendorf 5415 C centrifuge for 30 min at 14,000 rpm. The supernatant was removed and the pellet dried and resuspended in 14.5 μl DPBS buffer, 2.5 μl of DTT (10x), and 7 μl of LDS (4x).

### Gel electrophoresis and fluorography

All final samples were heated at 70 °C for 10 min in 1x of NuPAGE loading buffer (Invitrogen, Waltham, MA) before loading onto the NuPAGE gel (Invitrogen). Electrophoresis was performed using the NuPAGE system and precast Bis-Tris 10%, 12%, or 4-12% gradient gels and MES or MOPS running buffer. Gels were soaked in Amplify (Amersham, ArlingtonHeights, IL) or Enlightning (Revvity Health Sciences, Boston, MA) to enhance ^35^S fluorography, dried and exposed to Hyperfilm ECL film at –80°C. Typical exposure times were 16–30 h. Quantification of gels was carried out directly using a Molecular Dynamics PhosphorImager (Typhoon FLA 9500) and ImageQuant TL (v7.0) (Sunnyvale, CA).

### Molecular dynamics simulations of T1 domain

The intracellular domain of the structure of human Kv1.3 (PDB 7SSX^56^), determined by the single-particle cryoEM method, was used as the simulation template structure. Residues 103 through 206 in each subunit of the tetrameric complex was prepared in 150 mM KCl for all-atom molecular dynamics simulations using CHARMM-GUI^57^. A typical system consisted of 416 amino-acid residues in the four polypeptides, ∼12240 TIP3P waters, ∼38 K^+^ ions, and ∼34 Cl^−^ ions in a periodic boundary rectangular box of about 75 Å x 75 Å x 75 Å in size. The CHARMM36m all-atom force field formulation was utilized. The default equilibration parameter values suggested by CHARMM-GUI were used. The simulation runs at 90°C and 1 atm with Langevin piston were conducted with CUDA-enabled NAMD2 on x86-64-based computers running MacOS, Ubuntu or CentOS. The mutation R114G was introduced in each subunit with CHARMM-GUI during system preparation. Three independent WT systems and three independent R114G systems were prepared using CHARMM-GUI, and each system was simulated for 3 µs.

The trajectory results were analyzed every 1 ns using VMD (v1.9) and PyMol (v2.5 and v3.0, Schrödinger, New York, NY) on MacOS. Root-mean-square deviations (RMSDs) were calculated using the RMSD trajectory tool in VMD. In each simulated system, the simulated protein structures were first aligned to the starting structure and then non-hydrogen protein atom RMSDs were calculated. Atom displacements were also measured after alignment in PyMol. Nonbonded potential energy (Enb) levels of the simulated complexes were calculated with the NAMD energy plugin of VMD. Short-range nonbonded interactions were computed with a cutoff distance of 12 Å and with a switching function starting at 10 Å. Long-range electrostatic interactions were calculated with the particle mesh Ewald (PME) method with a grid spacing of 1.2 Å. The dihedral angles were measured in PyMol. Interaction energies were also calculated using the NAMD energy plugin of VMD utilizing the PME method as above. Some of the results were then exported to Igor Pro (v9, Wavemetrics, Lake Oswego, OR) and plotted.

### Structural visualization

Protein structures were visualized using RasMol (v. 2.7.5.2) running on Windows or PyMol (v.2.5.5) running on Windows.

## ACKNOWLEDGMENTS

We thank M. Bucan and A. Ghorai for useful conversations and pilot screening of KCNA3 PheCodes and S.Dudek for assistance with PheWAS View plots. The NIH Instrumentation Grant S10OD023592 Cluster was made available by K. Sharp, for molecular dynamics simulations. The work was supported by NIH grants GM52302 (C. Deutsch). R01GM121375 (T. Hoshi).

---

**Supplementary Table 1. Variant location and database parameters.** CADD PHRED scores (>20 indicates deleterious; PHRED scale scores are normalized to all potential 9 billion single nucleotide human variants) and Revel scores (rare missense variants >0.5 are likely disease-causing) for each variant. The UCSC Genome Browser was used to obtain Revel scores and the VarSite database (www.ebi.ac.uk/research/thornton/software/) was used to obtain CADD PHRED scores. The reference single nucleotide polymorphism (rs) ID in dbSNP (http://www.ncbi.nlm.nih.gov/SNP/) is a key resource identifier.

**Supplementary Figure S1. PheWAS-View plot for variants F203S** (**A.**) **and E182K** (**B.**) showing −log_10_ p value (left) and effect size estimates (beta; right) from PheWAS analyses on the x-axis and specific clinical phenotypes on the y-axis. This plot shows top 15 results from PheWAS analyses. Red line shows the line of significance (p value=.01). Gray line in S1B is a beta value of zero; values left of the gray line are negative beta values. AFR results are shown as blue points and EUR results are shown as red points. For broad phenotype categories, see Supplementary Figs. S2-S4.

**Supplementary Figure S2. Manhattan plot for variant R214C.** Manhattan plot for R214C showing the −log_10_ p values for all tested phenotypes across cohorts. The phenotypes are grouped by their disease categories on the x-axis. The black dashed line represents a significance threshold of .01.

**Supplementary Figure S3. Manhattan plot for variant F203S.** Manhattan plot for F203S showing the −log_10_ p values for all tested phenotypes across cohorts. The phenotypes are grouped by their disease categories on the x-axis. The black dashed line represents a significance threshold of .01.

**Supplementary Figure S4. Manhattan plot for variant E182K.** Manhattan plot for E182K showing the −log_10_ p values for all tested phenotypes across cohorts. The phenotypes are grouped by their disease categories on the x-axis. The black dashed line represents a significance threshold of .01.

**Supplementary Figure S5. Structural Models and nonbonded potential energies from multiple MD simulations. A.** Tetrameric structures derived from MD simulations for T1 WT and R114G at 1 ns and 3.0 μs viewed from the intracellular side. In each subunit (A, B, C, D), helices are colored red, β strands yellow, loops green. Rendered with PyMol. Symbols for three independent MD simulations of WT and of R114G are (O,Ñ, D) and (O,∇, Δ), respectively. **B.** Mean nonbonded potential energies (E_nb_) for individual subunits across three independent systems of WT (blue) and three independent R114G (red) tetrameric complexes. (n=12 subunits for each condition/curve.) Lightly shaded areas represent standard deviation.

