## Supplemental Table and Figures for "Disease-associated Kv1.3 variants are energy compromised with impaired nascent chain folding"

### Slide 1
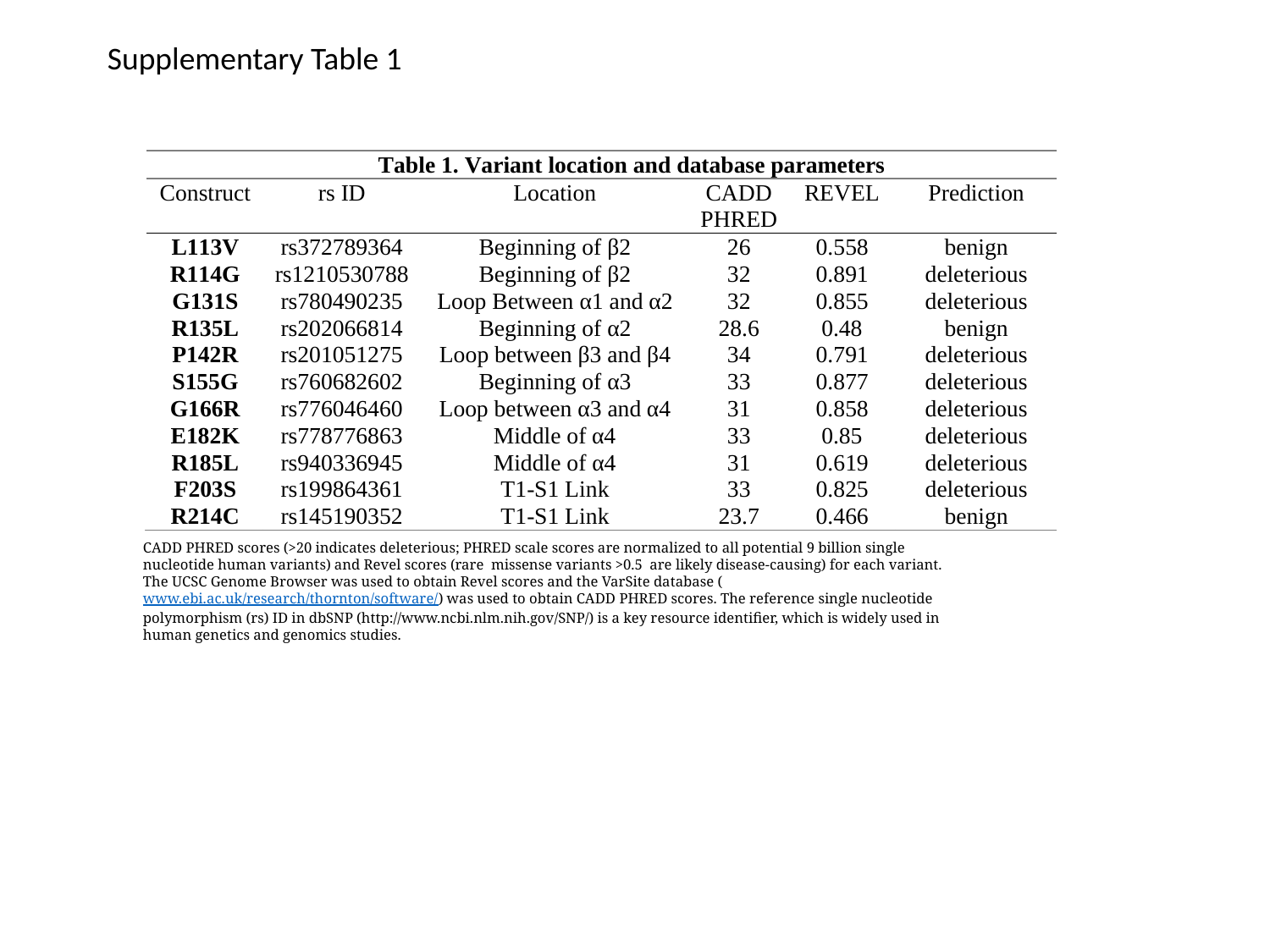

Supplementary Table 1
CADD PHRED scores (>20 indicates deleterious; PHRED scale scores are normalized to all potential 9 billion single nucleotide human variants) and Revel scores (rare  missense variants >0.5  are likely disease-causing) for each variant. The UCSC Genome Browser was used to obtain Revel scores and the VarSite database (www.ebi.ac.uk/research/thornton/software/) was used to obtain CADD PHRED scores. The reference single nucleotide polymorphism (rs) ID in dbSNP (http://www.ncbi.nlm.nih.gov/SNP/) is a key resource identifier, which is widely used in human genetics and genomics studies.

### Slide 2
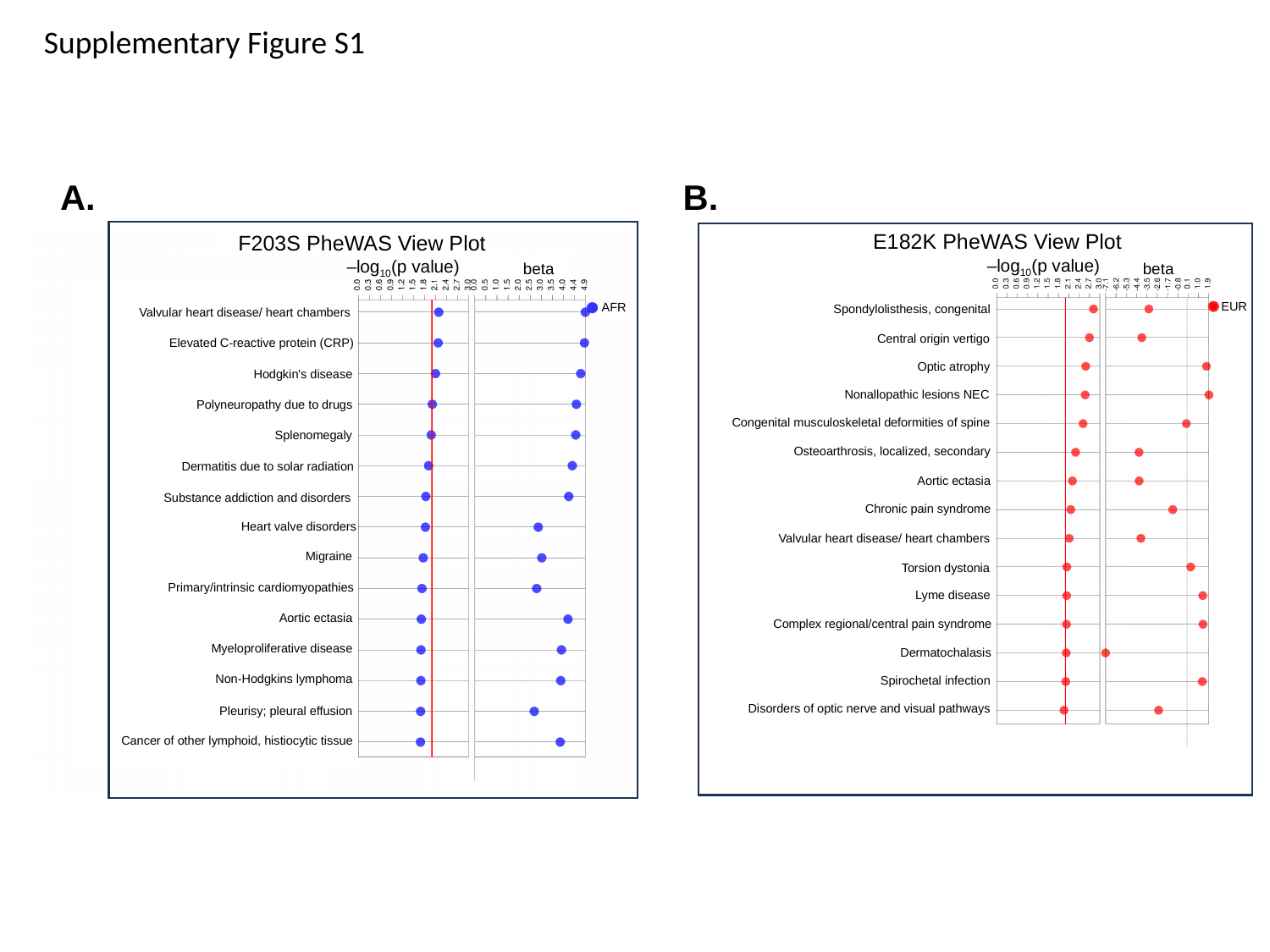

Supplementary Figure S1
A.
B.
F203S PheWAS View Plot
-log10(p value)
beta
AFR
Valvular heart disease/ heart chambers
Elevated C-reactive protein (CRP)
Hodgkin's disease
Polyneuropathy due to drugs
Splenomegaly
Dermatitis due to solar radiation
Substance addiction and disorders
Heart valve disorders
Migraine
Primary/intrinsic cardiomyopathies
Aortic ectasia
Myeloproliferative disease
Non-Hodgkins lymphoma
Pleurisy; pleural effusion
Cancer of other lymphoid, histiocytic tissue
‒log_10_(p value)
E182K PheWAS View Plot
-log10(p value)
beta
EUR
Spondylolisthesis, congenital
Central origin vertigo
Optic atrophy
Nonallopathic lesions NEC
Congenital musculoskeletal deformities of spine
Osteoarthrosis, localized, secondary
Aortic ectasia
Chronic pain syndrome
Valvular heart disease/ heart chambers
Torsion dystonia
Lyme disease
Complex regional/central pain syndrome
Dermatochalasis
Spirochetal infection
Disorders of optic nerve and visual pathways
‒log_10_(p value)
PheWas View Plot for F203S

### Slide 3
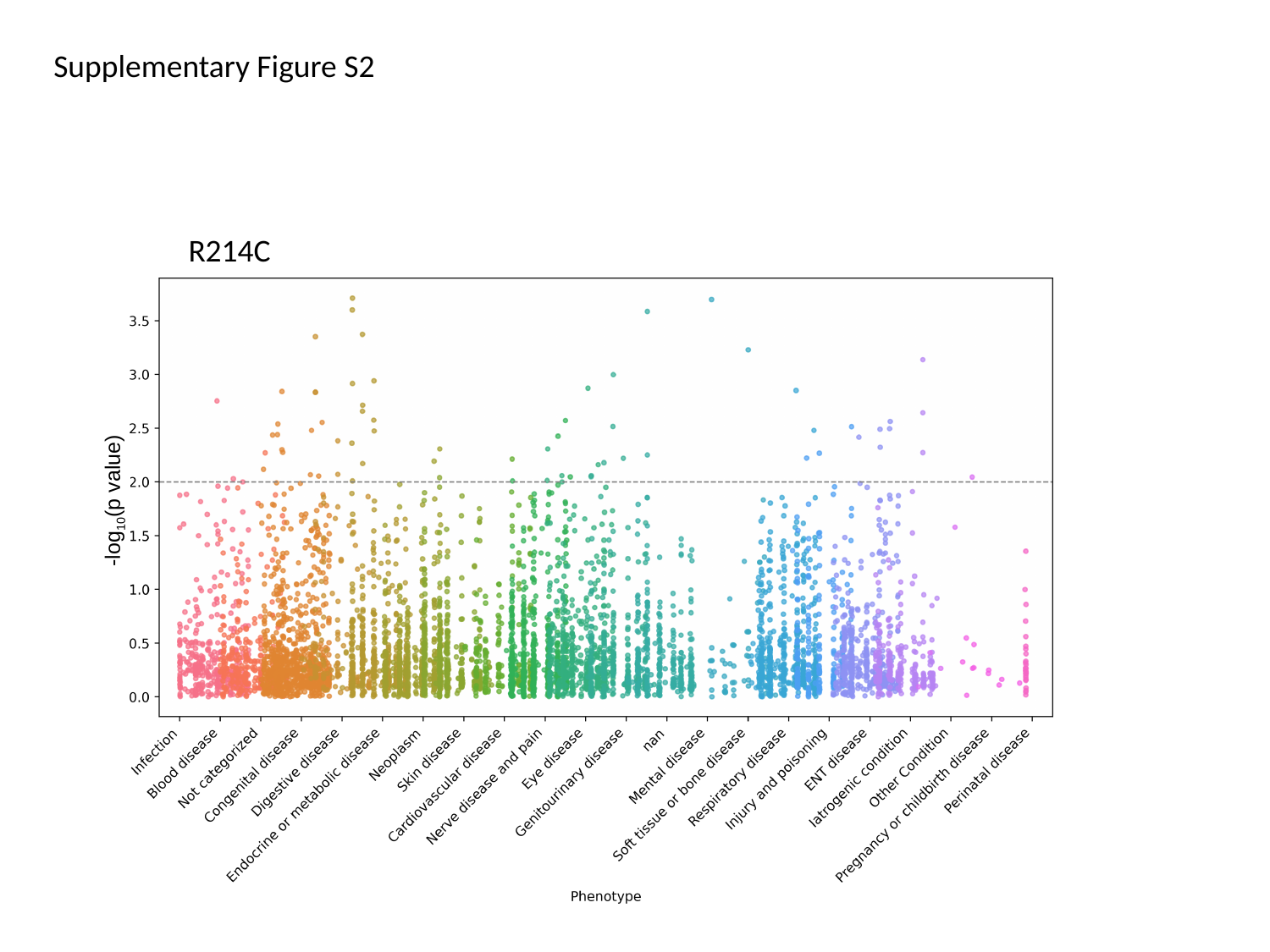

Supplementary Figure S2
-log_10_(p value)
R214C

### Slide 4
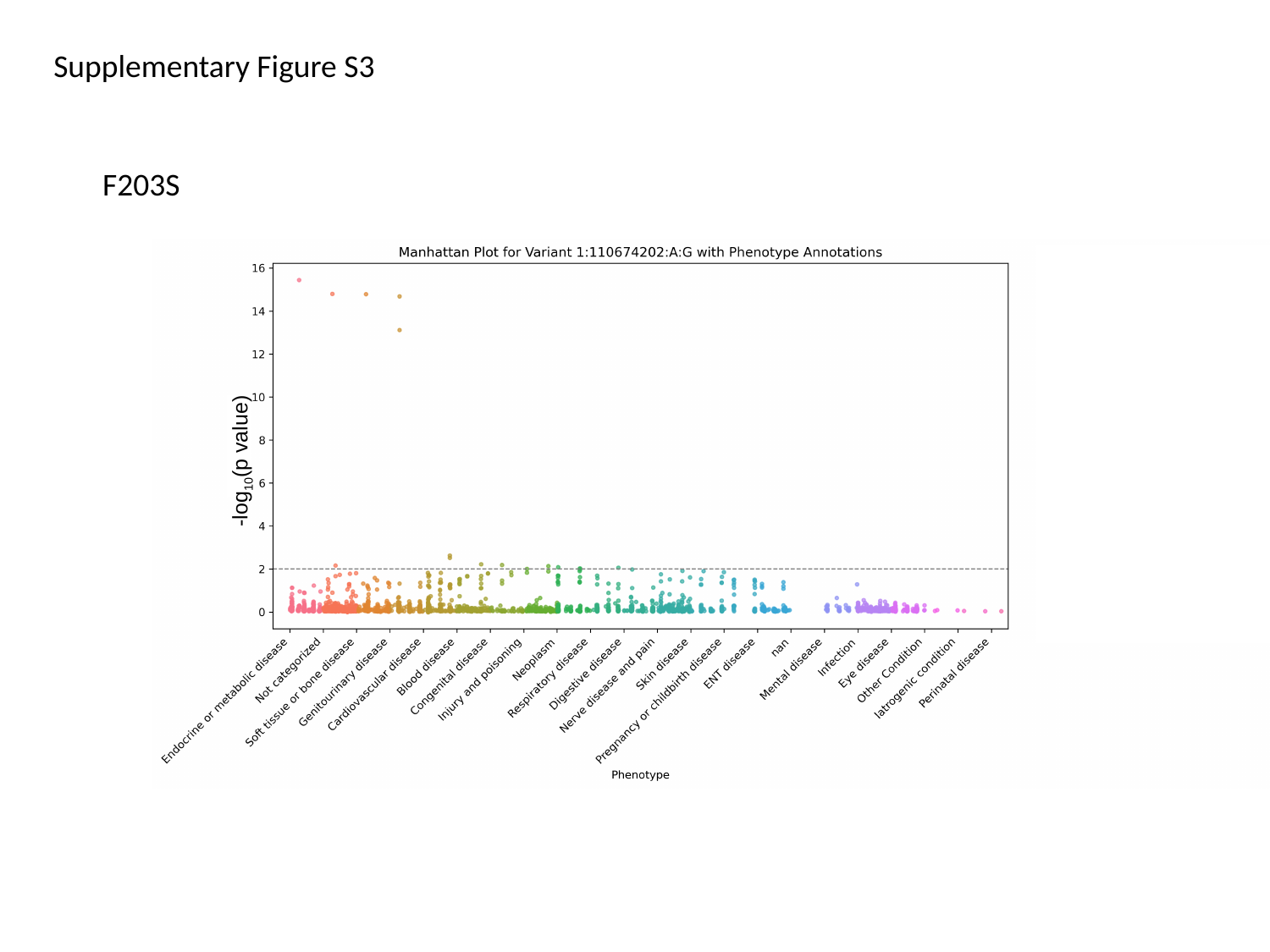

Supplementary Figure S3
F203S
-log_10_(p value)

### Slide 5
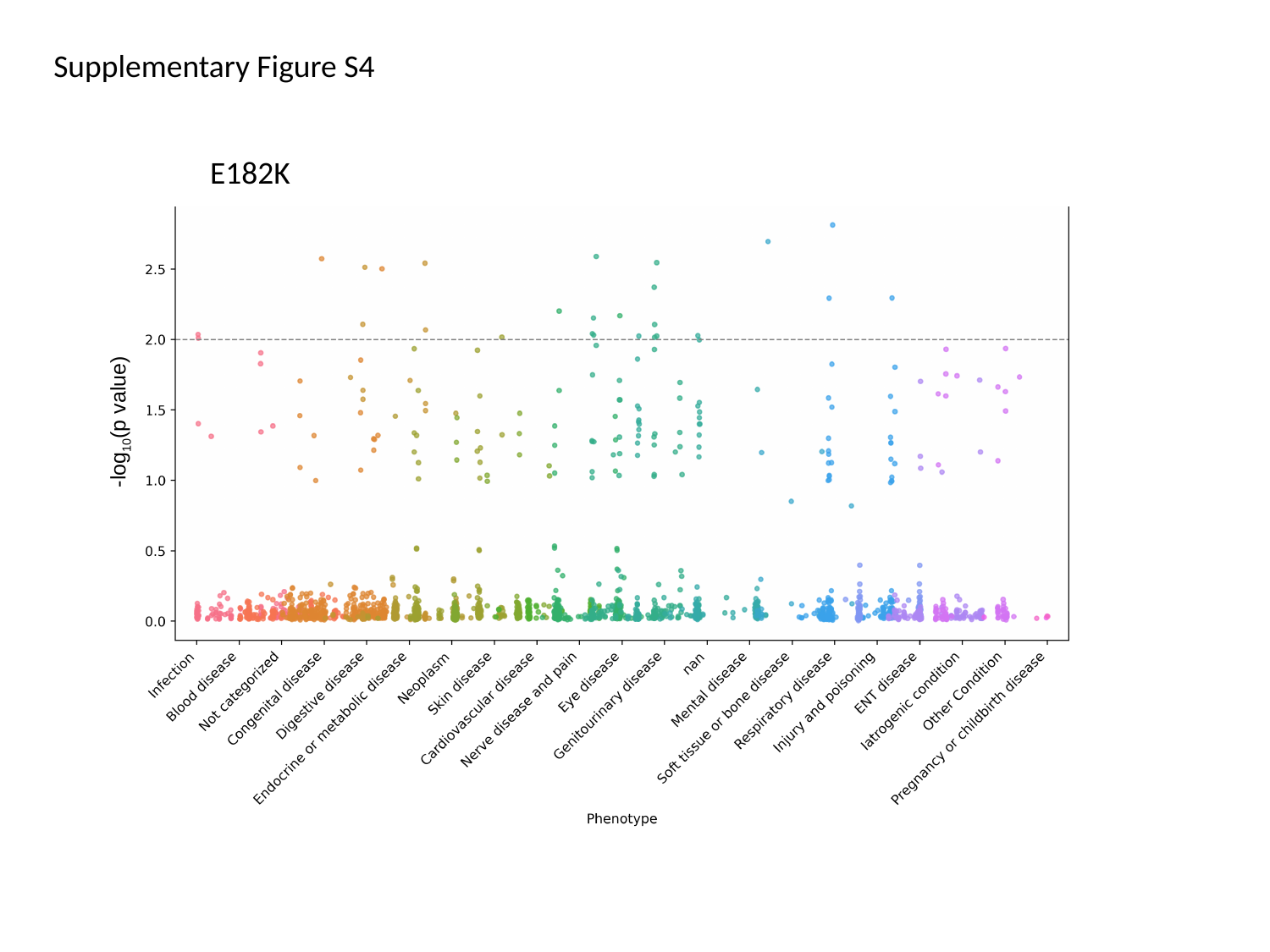

Supplementary Figure S4
E182K
-log_10_(p value)

### Slide 6
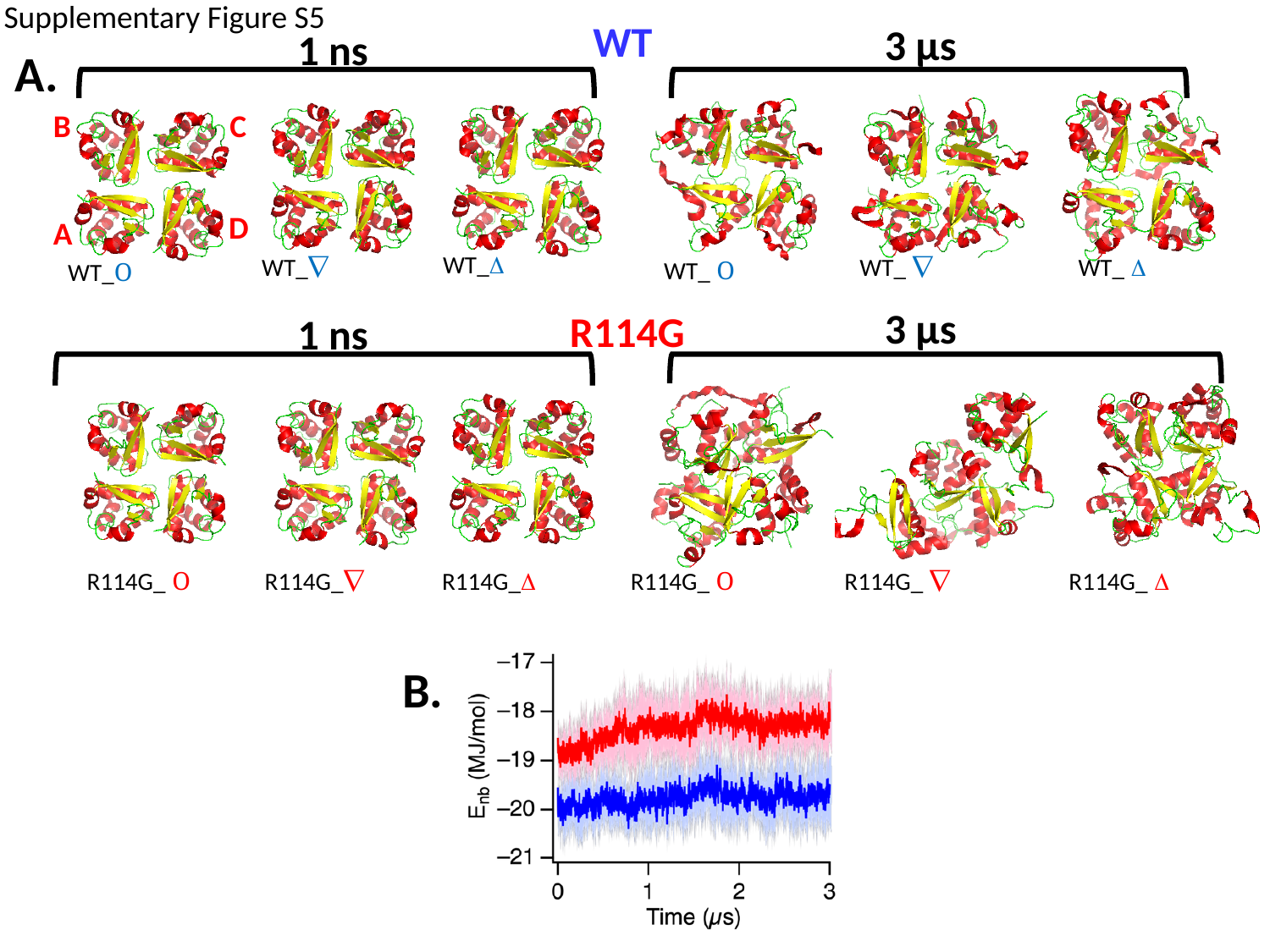

Supplementary Figure S5
WT
3 μs
1 ns
A.
B
C
D
A
WT_Δ
WT_∇
WT_ ∇
WT_ Δ
WT_ O
WT_O
3 μs
R114G
1 ns
R114G_ O
R114G_∇
R114G_Δ
R114G_ O
R114G_ ∇
R114G_ Δ
B.
